# Crystal structure of m4-1BB/4-1BBL complex reveals an unusual dimeric ligand that undergoes structural changes upon receptor binding

**DOI:** 10.1101/444554

**Authors:** Aruna Bitra, Tzanko Doukov, Giuseppe Destito, Michael Croft, Dirk M. Zajonc

## Abstract

The interaction between the 4-1BB and its ligand 4-1BBL provides co-stimulatory signals for T cell activation and proliferation, but differences in the mouse and human molecules might result in differential engagement of this pathway. Here, we report the crystal structure of mouse 4-1BBL and of the mouse 4-1BB/4-1BBL complex, together provide insights into the molecular recognition of the cognate receptor by m4-1BBL. In contrast to all human or mouse TNF ligands that form non-covalent mostly trimeric assemblies, the m4-1BBL structure formed a novel disulfide linked dimeric assembly. The structure showed that certain differences in the amino acid composition along the intramolecular interface, together with two specific residues (Cys 246 and Ser 256) that are exclusively present in m4-1BBL, are responsible for unique dimerization. Unexpectedly, upon binding to m4-1BB, m4-1BBL undergoes structural changes within each protomer, in addition the individual m4-1BBL protomers rotate with respect to each other, leading to a different dimerization interface with more inter-subunit interactions. In the m4-1BB/4-1BBL complex, each receptor monomer binds exclusively to a single ligand subunit with contributions of cysteine-rich domain (CRD) 1, CRD2 and CRD3. Furthermore, structure-guided mutagenesis of the binding interface revealed that novel binding interactions with the GH loop, rather than the DE loop, are energetically critical and define the species based receptor selectivity for m4-1BBL. A comparison with the human 4-1BB/4-1BBL complex highlighted several differences between the ligand and receptor binding interfaces and provide an explanation for the absence of inter species cross-reactivity between human and mouse 4-1BB and 4-1BBL molecules.

---

4-1BB is a type 1 trans membrane protein of the tumor necrosis factor receptor superfamily (TNFRSF) that is expressed on multiple cell types including T cells, dendritic cells and NK cells (1,2). The ligand 4-1BBL is a type II trans membrane protein expressed on antigen presenting cells, such as B cells, macrophages and dendritic cells and its expression is upregulated upon stimulation (3,4). Similar to other TNFRSF members, aggregation of 4-1BB via binding to its ligand results in the recruitment of intracellular TRAF adaptor molecules (TRAF1 and TRAF2), leading to activation of several proinflammatory-signaling pathways (1,5,6). Binding of 4-1BBL to 4-1BB generates strong co-stimulatory signals in T cells that lead to upregulation of anti-apoptotic molecules, cytokine secretion, and enhanced effector function (7,8).

Most of the members of the TNFR family are monomeric and share a similar topology composed of several cysteine rich domains (CRDs) within the extracellular domain and TRAF binding motifs in their cytoplasmic regions (9). A typical CRD is composed of cysteine residues that form intra-molecular (and intra-domain) disulfide bonds and according to their number and topological connectivity, each CRD can contain any of three modules A1, A2 and B2 (10). The ectodomain of human and mouse 4-1BB contains 10 intra-disulfide bonds that maintain the structural and functional integrity of the protein. Human 4-1BB contains an additional cysteine at position 121 in CRD4 that forms an inter-molecular disulfide bond between two adjacent monomers (11). The CRD1 region of both human and mouse 4-1BB is partial and it lacks a conserved anti parallel β strand motif. The CRD2 contains A1 and B2 modules made up of antiparallel β strands with 1-3 and 2-4 disulfide connectivity and the CRD3 contains A1 and A2 modules. Though both 4-1BB molecules exhibit ~ 30% sequence identity with other characterized TNFR members, the CRD3 and CRD4 regions do not superimpose with any other receptor. This is due to the bend in the central hinge region of both human and mouse 4-1BB that joins CRD2 and CRD3, which changes the relative orientation of CRD3 and CRD4 distinctly with respect to corresponding regions of other TNFR members (12).

All of the human TNF family ligands are trimeric, and display a classic THD β sandwich jellyroll fold (13,14). However, they exhibit structural diversity in the way the individual subunits assemble with respect to each other. In this regard, the TNF ligands were originally described to fall into three sub-families based on their sequence variance and structural organization (14). The majority of molecules are considered as conventional family ligands and contain longer loops connecting CD, DE and EF strands and are assembled as compact bell shaped trimers (15,16). In contrast, the EF-disulfide family ligands (APRIL, TWEAK, BAFF and EDA) are more globular because of shorter CD and EF loops and they bind to very small atypical TNFRSF members (13,17). While the conventional members possess a disulfide bond linking CD and EF loops, the EF-family members contain a disulfide bond linking E and F strands (14). The divergent family members are unique as they exhibit very low sequence similarity with other TNF ligands. The members of this family, OX40L and GITRL, possess shorter THD regions and assemble as more planar blooming flower shaped trimers (18,19). Based on sequence diversity, human 4-1BBL was previously categorized within the latter group. However, two recent crystal structures of h4-1BBL revealed that h4-1BBL forms a compact bell shaped trimer characteristic of conventional TNF ligands (11,20). All conventional ligands bind to their cognate receptors in a similar manner and they all contains a conserved hydrophobic residue that acts as a ‘hot spot’ in their DE loop that is shown to be energetically important for receptor binding (13). However, the crystal structure of OX40/OX40L revealed that the binding energy is distributed equally on both sides of the interaction interface and there is no significant hydrophobic contact between the DE loop of the ligand and the receptor (18). Interestingly, although human 4-1BBL exhibits all the features of the conventional ligands, the hydrophobic residues in the DE loop are not conserved. Additionally, recent crystal structures showed that the DE loop residues are not contributing towards binding affinity, suggesting that h4-1BB although forming a bell-shaped trimer is still unique in its interaction with h4-1BBL (11,20,21).

While many TNF-TNFR complexes had been crystallized, until recently, very little was known about 4-1BB/4-1BBL interactions. Our recent characterization of the interaction between human 4-1BB and 4-1BBL led us to hypothesize a distinct mechanism of h4-1BB signaling, in which covalent receptor-dimerization would favor the formation of a 2D-signaling network to initiate robust signaling (11,20). Adding to this, our earlier biochemical studies suggested that recombinant m4-1BBL could form a covalent dimer rather than the conventional trimer (12). This supports a different mechanism of human and mouse 4-1BBL to both engage and cluster 4-1BB, using unique protein binding sites and resulting in different oligomeric assemblies that would affect the signal strength. To elucidate the molecular features underlying the differential behavior of m4-1BB and m4-1BBL, we have determined the crystal structure of m4-1BBL itself as well as in complex with its receptor m4-1BB. Together, our findings identify structural details related to ligand-receptor interactions but also the assembly of the unique m4-1BBL.

## RESULTS

### Crystal structure of m4-1BB ligand ectodomain

Mouse 4-1BBL (m4-1BBL) is a type II transmembrane protein composed of an N-terminal cytoplasmic region and a C-terminal ectodomain separated by a transmembrane domain. The ectodomain can be divided into a tail region and the TNF homology domain (THD) (Figure 1A). The THD is responsible for the interaction with its cognate receptor m4-1BB. Two N-linked glycosylation sites are present, one at the N-terminal (Asn 161) and the other at the C-terminal end (Asn 293) of the THD (Fig.1A). We crystallized the THD of m4-1BBL along with several additional C-terminal tail residues (amino acids 140-309) and determined the structure by molecular replacement using h4-1BBL (PDB ID 6D3N) as a search model (Table 1). The asymmetric unit in the crystal contained four copies of m4-1BBL and in the final model, the amino acids at the N- and C-terminal ends and some of the surface exposed loops were disordered due to their high flexibility. In all four copies, we were able to build N-glycans at Asn 161; however, we have not observed obvious electron density for glycans at Asn 293. The global superposition of all copies of m4-1BBL in the asymmetric unit indicates high similarity in their structure with marginal variation in their loop regions (root mean square deviation value (rmsd) of 0.124 Å) and conserved orientation of the N-glycans.

**FIGURE 1.**
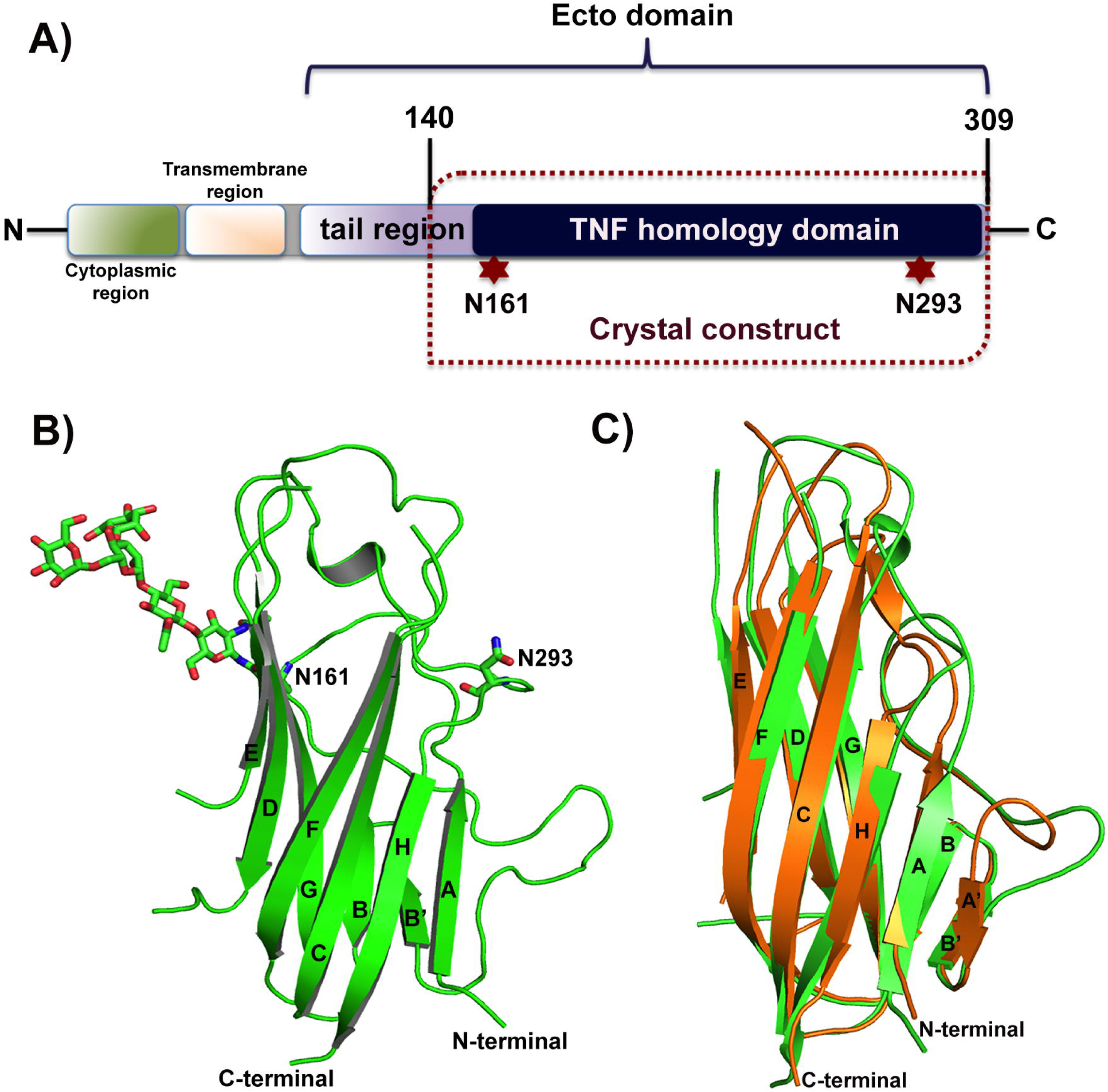
Architecture of m4-1BBL. A) Domain organization of m4-1BB ligand. The construct used for crystallization is highlighted in dashed lines. Two N-linked glycosylation sites at positions 161 and 293 are highlighted. B) Cartoon representation of m4-1BBL monomeric structure. The β strands are labeled in accordance to the structure of h4-1BBL (PDB 6D3N) and the N/C-terminal ends are marked. Asparagine residues of the potential N-glycosylation sites and N-glycans at Asn 161 position are shown as sticks. C) Structural superposition of monomeric m4-1BBL (green) and h4-1BBL (orange) illustrates differences along the β strands and the surface loops that connect these strands.

**Table 1.** Data collection and refinement statistics.

| Data collection statistics | M4-1BB ligand | M4-1BB/4-1BBL complex |
| --- | --- | --- |
| PDB ID | 6MKB | 6MKZ |
| Space group | P3<sub>1</sub> | P 2<sub>1</sub> 2<sub>1</sub> 2<sub>1</sub> |
| Cell dimension |  |  |
| *a*, *b*, *c*, (Å) | 77.3, 77.3, 118.2 | 61.5, 88.4, 155.1 |
| $\alpha$, $\beta$, $\gamma$(°) | 90.00, 90.00, 120.00 | 90.00, 90.00, 90.00 |
| Resolution range (Å) | 50.0 – 2.50 | 76.9 – 2.62 |
| [outer shell] | [2.59-2.50] | 2.69-2.62] |
| No. of unique reflections | 27191 [2727] | 25447 [1263] |
| R<sub>meas</sub> (%) | 8.1 (83.2) | 24.0 (101.2) |
| R<sub>pim</sub> (%) | 2.5 [25.6] | 6.7 [38.1] |
| Multiplicity | 10.4 (10.4) | 13.7 (7.5) |
| Average I/σ | 23 [2.0] | 5.6 [1.7] |
| Completeness (%) | 99.7 [99.9] | 97.6 [71.0] |
| Refinement statistics |  |  |
| No. atoms | 5706 | 4202 |
| Protein | 5175 | 3926 |
| Water | 286 | 198 |
| Glycerol/Na/sulfate/N-glycans | 245 | 78 |
| Ramachandran plot (%) |  |  |
| Favored | 95.9 | 94.4 |
| Allowed | 4.0 | 5.1 |
| Outliers | 0.2 | 0.6 |
| R.m.s. deviations |  |  |
| Bonds (Å) | 0.009 | 0.010 |
| Angles (°) | 1.18 | 1.24 |
| B-factors (Å<sup>2</sup>) |  |  |
| Protein | 56.3 | 60.5 |
| Water | 58.1 | 61.4 |
| Glycerol/Na/Cl/sulfate/N-glycans | 64.6 | 53.7 |
| R factor (%) | 18.8 | 20.4 |
| R<sub>free</sub> (%) | 24.1 | 23.8 |

Each m4-1BBL monomer adopts a canonical β-sandwich jelly-roll fold composed of two anti-parallel inner and outer β-sheets formed by AHCF strands and B’BGDE strands respectively (Figure 1B). The N-terminal A’ strand present in other human or mouse TNF ligands is substituted by the longer AB’ loop in m4-1BBL. The THD of m4-1BBL shares modest sequence identity (~ 40%) with its human homologue, h4-1BBL, but they both display considerable topological similarity at their β-strand regions (with an RMSD of 0.8 Å on equivalent CA atoms) (Figure 1C). The only pronounced differences between the monomeric m4-1BBL and h4-1BBL structures are present in the conformations of the side chains and the surface loops that connect the β-strands. The length of the THD of m4-1BBL is ~ 55 Å, which is comparable to that found in canonical TNFs like RANKL, CD154, and others. In addition, m4-1BBL also contains extended C, D, E and F strands and elongated loops connecting these strands. In summary, the protomer of both m4-1BBL and h4-1BBL are structurally very similar and also share structural details with members of the conventional TNF family.

### Unique dimeric organization of m4-1BB ligand

To date, all the characterized conventional human or mouse TNF ligands organize into a symmetrical trimeric bell shape to form a functional biological unit. Recently, we and others reported the structure of h4-1BBL that also assembles as a trimeric bell shape despite its low sequence similarity with conventional members (11,20). However, m4-1BBL self-assembled as a two-fold symmetrical homodimer, in which both protomers are covalently connected by a disulfide bond (Figure 2A). The dimeric interface is formed by inner sheet β strands of both protomers that pack against each other with a total buried surface area of 1617 Å^2^. At the top of the dimer, the EF loop of both protomers are placed nearby, and as a consequence, their surface exposed cysteine residues (Cys 246) interact to form an inter-molecular disulfide linkage between them. Unexpectedly, other than the disulfide bond between two protomers, the m4-1BBL dimer lacks any intersubunit contacts at the upper and middle half of the protomers. But, at the lower half of the dimer, the residues Tyr 199, Phe 201, Phe 300, Val 302 and Phe 148 from both protomers form a hydrophobic core to mediate strong stabilizing interactions (Figure 2B). In addition, at the lower end, the N-terminal Pro 146 of one protomer interacts with the C-terminal Pro 304 of the second protomer via van der Waals contacts, thereby closing the tunnel-like opening that is running downward from the top of the dimer. Throughout the structure, we have not seen any potential hydrogen bonds at the monomer-monomer interface while most of the TNF ligands possess around 10-15 hydrogen bonding interactions promoting stabilization of the trimeric interface.

**FIGURE 2.**
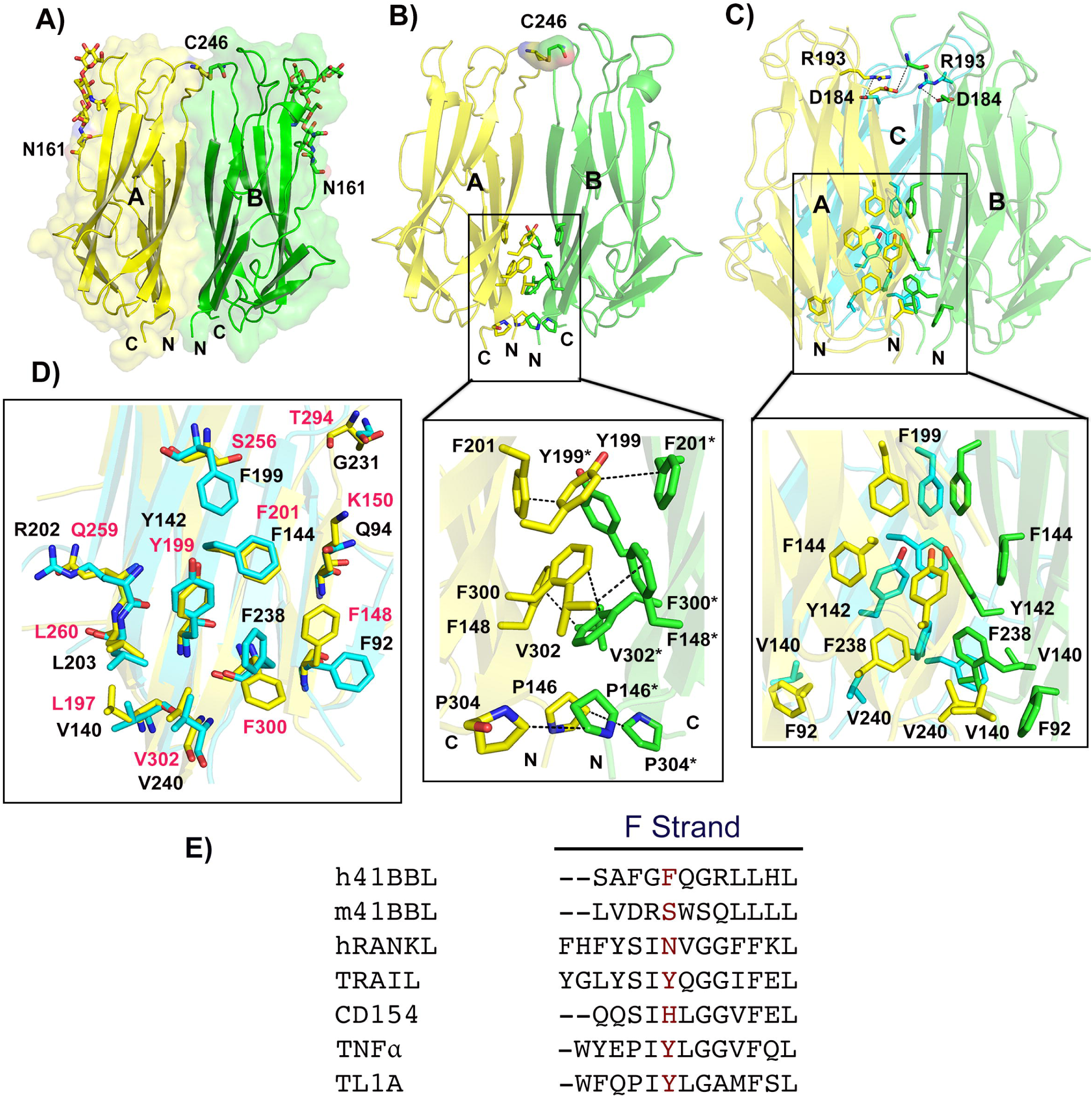
Structure of m4-1BBL dimer. A) Cartoon representation of m4-1BBL dimer with transparent molecular surface. The N-terminal and C-terminal ends of both protomers are marked. The N-glycans at Asn 161 position are shown as ball and sticks. B) Dimerization interface of m4-1BBL dimer. All residues mediating hydrophobic interactions at the lower interior of the dimer are shown as sticks in the inset below. In Panel A and B, the carbon atoms of residues of protomer A are shown in yellow color and protomer B are in green. In both panels, the cysteine residues (Cys 246) that form disulfide linkage between two protomers at the top of the dimer are shown as sticks. C) Trimerization interface of h4-1BBL. The three protomers of h4-1BBL trimer are colored yellow, green and cyan. The residues mediating salt bridge interactions and vdW’s contacts among all three protomers of h4-1BBL are labeled. D) Structural alignment of residues mediating stabilization of monomer-monomer interface in h4-1BBL and m4-1BBL. The residues of m4-1BBL are shown in yellow with red labels and residues of h4-1BBL are shown in cyan with black labels. E) Structure based sequence alignment of ‘F’ strand residues of conventional TNF ligands. In each ligand, the residue corresponding to Phe 199 of h4-1BBL is colored in red.

Although the individual protomers of mouse and human 4-1BBL are largely comparable, superposition of dimeric m4-1BBL with the corresponding regions on the h4-1BBL trimer results in higher RMSD values due to the variation in the alignment of its individual protomers with respect to each other (Figure S1A). In h4-1BBL, the trimer is arranged such that the E and F strand of all three protomers packed against the inner sheet of its adjacent protomer, thus hydrophobic and polar residues from inner β strands A, C, E, F and H mediate stabilization of the trimer (Figure 2C and Figure S1B). In addition, the surface loop connecting the strands E and F also contribute residues that mediate potential hydrogen bonding or salt bridge interactions in the trimeric interface (11) (Figure 2C). But, owing to the disparity in subunit orientation, only hydrophobic residues from A, C and H strands form the dimeric interface in m4-1BBL (Figure 2B and Figure S1C). None of the E and F strand residues or the surface loops partake in mediating interactions between two subunits, and so the top and central portion of the dimer lacks intersubunit contacts. As a consequence, the contact surface area between m4-1BBL protomers is considerably reduced with the interfacial organization formed by a total of ~ 20 residues while in h4-1BBL many more residues (~ 40) are involved in trimer interactions.

A structure-based sequence alignment of both mouse and human 4-1BB shows that most of the residues mediating trimerization in h4-1BBL are present in m4-1BBL (Figure 2D). Also, at the lower half of the monomer-monomer interface, all of those residues involved in the hydrophobic core are substantially similar and ~ 80% of them are even conserved in both species (Figure 2D). However, at the middle half of the dimer, m4-1BBL contains a hydrophilic residue (Ser 256) on the F strand while h4-1BBL features a phenylalanine residue (F199) at this position (Figure 2D). The structure of trimeric h4-1BBL reveals that the aromatic side chain of Phe 199 from all three subunits protrudes towards the interior to form a hydrophobic triade (Figure 2C). Structural alignment with other TNF ligands demonstrates that this hydrophobic core located on the ‘F’ strand is conserved in hTRAIL, hTNF and hTL1A (either phenylalanine or tyrosine residue) but not in m4-1BBL, hRANKL and hCD40L (Figure 2E). In the absence of hydrophobic residues, the corresponding amino acid facilitates polar contacts between protomers. For instance, in hRANKL, the equivalent Asn 276 from all three protomers interact via hydrogen bonding to promote intersubunit contacts, while similar type of interactions are not observed in m4-1BBL (using Ser 256).

These data suggest that although m4-1BBL contains most of the characteristic features required for packing into a bell-shaped trimer, the lack of stabilizing interactions from the E / F strands and the unique EF loop may interfere with trimerization and instead result in an atypical dimeric structure. Therefore, to explore the contribution of Cys 246 and Ser 256 in mediating the dimerization of m4-1BBL, we sequentially mutated the Cys 246 to serine and Ser 256 to phenylalanine. In addition, we also made a double mutant of m4-1BBL carrying both of these mutations. The SDS PAGE analysis of all of these variants revealed that under non-reducing conditions, the C246S and C246S/S256F mutants of m4-1BBL migrate as a double band with equal intensity corresponding to the monomer size of ~26 kDa whereas the S256F mutant migrates similar to dimeric wildtype m4-1BBL of ~56 kDa molecular weight (Figure 3A). As expected, removal of Cys 246 abrogates the covalent dimerization. However, the S256F variant with Cys 246 intact still forms a covalent dimer, suggesting that the S256F mutation is not sufficient to prevent the formation of the intermolecular disulfide bond. Since the m4-1BBL subunit has a calculated molecular weight of ~20 kDa plus additional mass from N-glycans, treatment of the m4-1BBL monomer variants with PNGaseF now results in m4-1BBL migrating at a single band of ~20 kDa suggesting that the double band was due to differential glycosylation of both m4-1BBL protomers.

**FIGURE 3.**
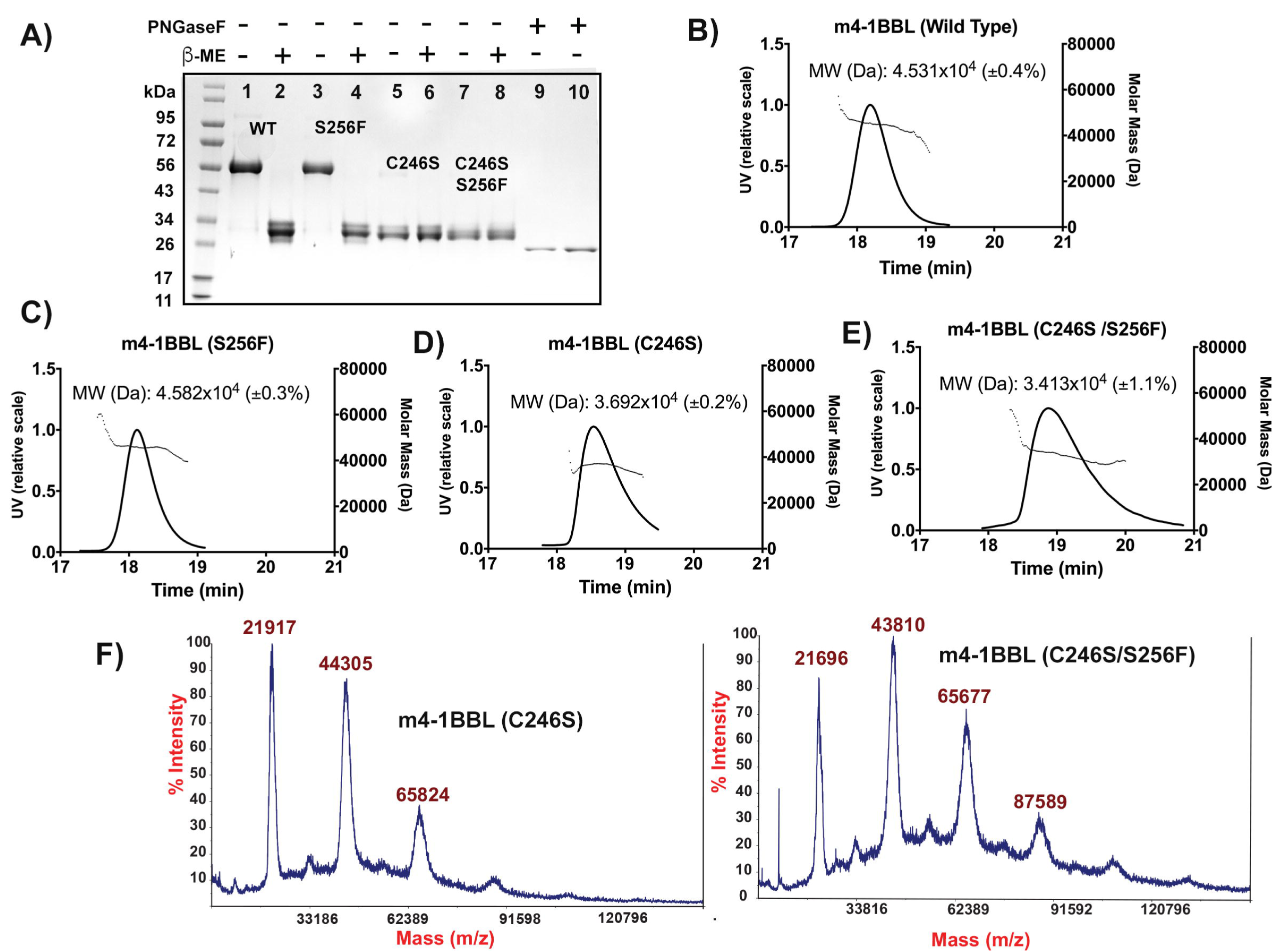
Characterization of m4-1BBL dimerization mutants. A) The SDS PAGE (4-20%) analysis of purified WT and various mutants of m4-1BBL under non-reducing (-β-ME (Lane 1, 3, 5, 7) and reducing conditions (+β-ME (Lane 2, 4, 6, 8). Lane 1and 2 corresponds to m4-1BBL (WT); lane 3 and 4 corresponds to m4-1BBL (S256F); lane 5, 6 corresponds to m4-1BBL (C246S); lane 7, 8 corresponds to m4-1BBL (C246S/S256F); lane 9 and 10 represent deglycosylated (+PNGase) WT and C246S mutant of m4-1BBL. B-E) SEC-MALS analysis of wildtype, S256F, C246S and C246S/S256F variants of m4-1BBL. The molecular mass calculated from MALS data are indicated. F) Mass spectrometry analysis of C246S and C246S/S256F variants of m4-1BBL. Peaks are labeled with corresponding molecular weight in Da.

To verify the oligomeric state of all of these variants in solution, we performed SEC-MALS (Size exclusion chromatography combined with multi angle light scattering) studies. On a standardized HPLC column, the S256F mutant of m4-1BBL eluted as a sharp monodisperse peak, which is comparable to that of 44 kDa reference peak (Figure 3B and 3C). The C246S and C246S/S256F mutants of m4-1BBL ran smaller and a clear peak shift was observed in relation to the wildtype m4-1BBL. However, rather than adopting a molecular weight corresponding to a monomeric state, the MALS study calculated the molecular mass for both of these mutants to be approximately 35 kDa (Figure 3D and 3E). This suggests that even without the inter-molecular disulfide bond, m4-1BBL still forms a dimer but adopts a more compact shape that allows the protein to migrate faster on SEC and appear smaller using MALS. Compared to the C246S mutant, the effect of the C246S/S256F double mutant is more complex. Firstly, the SEC peak is broader compared to the C246S mutant, which is also reflected by the discrepancy in the molar mass across the peak as assessed by MALS. This could be because of the equilibrium shift from a potential dimer-monomer or dimer-trimer that occurs during the chromatographic run. A similar phenomenon has been reported for hemoglobin, which also exists in tetramer-dimer equilibrium during SEC-MALS analysis with a calculated averaged molecular weight (22). To further identify the different possible oligomeric states of the m4-1BBL mutants, we performed MALDI-TOF analysis. The C246S/S256F double mutant revealed three peaks of almost comparable intensity corresponding to a monomeric, dimeric, and trimeric arrangement of m4-1BBL, while the C246S single mutant has a trimeric peak of much lower intensity compared to the peaks correlating with the monomeric or dimeric form (Figure 3F). This suggests that both C246 and the S256 are involved in the unusual dimeric assembly of m4-1BBL and that they may allow transition between monomeric, dimeric and trimeric arrangements, correlating with the broad peak obtained during SEC-MALS (Figure 3E).

### Interactions between m4-1BB ligand and its cognate receptor m4-1BB

As the disulfide linked m4-1BB ligand dimer is unique among TNF family members, we next determined its interaction with its cognate receptor m4-1BB using X-ray crystallography. We determined the crystal structure of the m4-1BB/4-1BBL complex at a resolution of 2.65 Å (Table 1). The asymmetric unit of the crystal contains two copies of the complex. Each protomer of the disulfide linked m4-1BBL binds one monomeric m4-1BB receptor, leading to a 2:2 arrangement as the minimal biological unit. In the final structure, with the exception of some flexible loops, most of the m4-1BBL structure was well ordered in both protomers (amino acids 145-308). However, while ~90% (amino acids (24-136; 151-155) of one of the two monomeric m4-1BB receptors was ordered, only ~70% of the second receptor was ordered. This may be due to crystal packing. Investigation of the m4-1BB/4-1BBL interaction reveal several structural features that were not previously witnessed in any of the TNF-TNFR complexes. In the complex, a single m4-1BBL engages a monomeric receptor leading to the formation of a tetrameric ligand-receptor complex. In all other reported TNF-TNFR complexes, including the recently solved structure of the human 4-1BBL/4-1BB complex (11,20), each receptor binds between two adjacent ligand protomers. M4-1BB, however, exclusively binds to one ligand protomer (protomer A) with no obvious contact with the adjacent subunit (protomer B) (Figure 4A). The high affinity interaction between m4-1BB and its ligand is evident from the extensive interface area of the complex in which ~1040 Å^2^ area is buried on the ligand and 990 Å^2^ is buried on the receptor.

**FIGURE 4.**
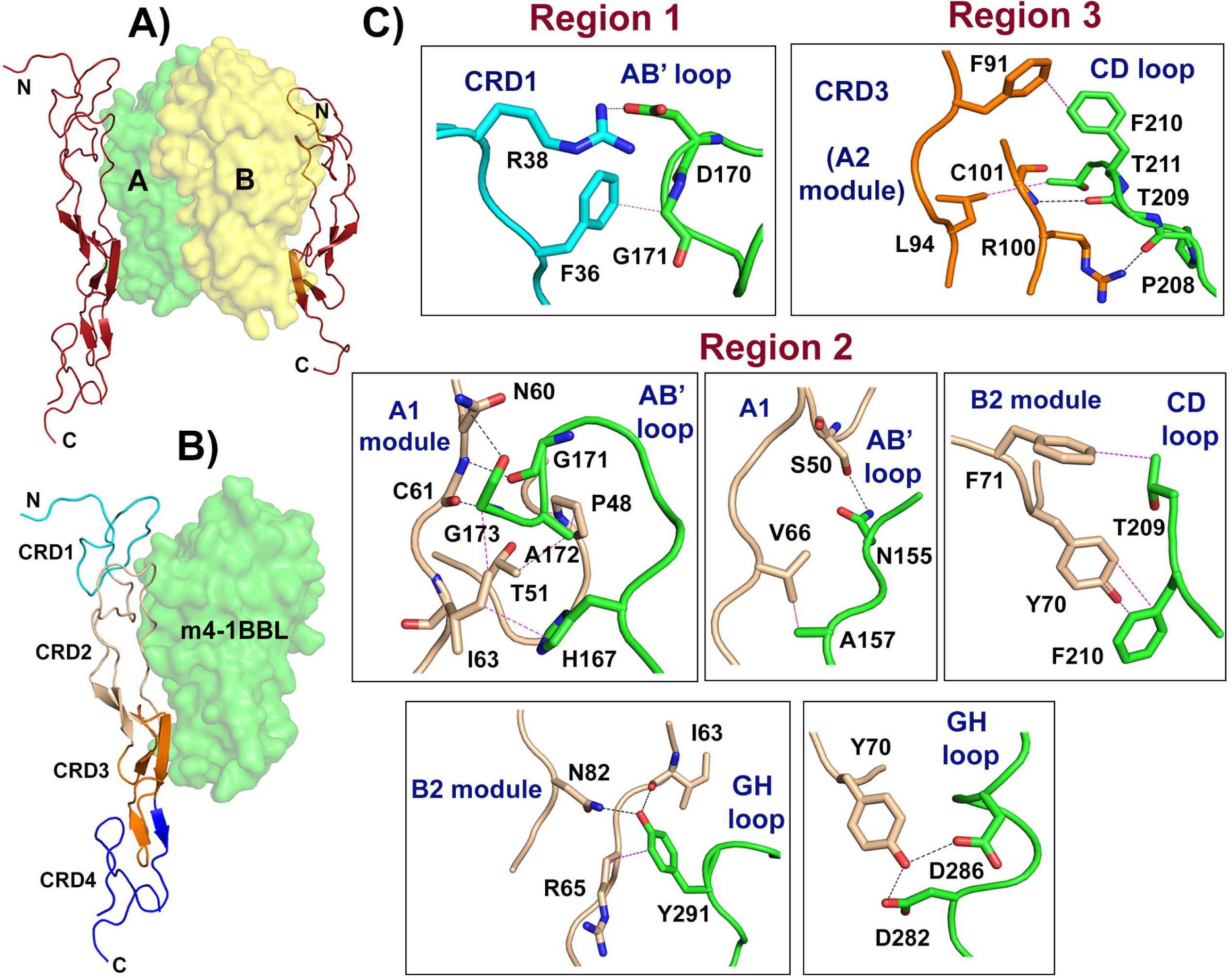
Structure of m4-1BB/4-1BBL complex. A) Crystal structure of tetrameric m4-1BB/4-1BBL complex. Two protomers of m4-1BBL are shown as transparent surface and the m4-1BB receptors are shown as cartoon. The N-terminal and C-terminal ends of receptor molecules are marked. B) Binding interface between m4-1BBL and m4-1BB. The ligand is shown as green transparent surface. The four CRDs of m4-1BB are colored individually and labeled. C) Detailed view of interactions between m4-1BBL and m4-1BB. The residues of m4-1BBL involved in the binding interface are shown as sticks in green color and their respective loops are labeled. The residues of m4-1BB are colored according to their corresponding CRDs; CRD1 as cyan, CRD2 as light brown and CRD3 as orange color. The different types of modules in CRD2 and CRD3 are labeled. The hydrogen bonding interactions are shown as black dashed lines and the hydrophobic contacts as magenta dashed lines.

M4-1BB predominantly uses its CRD2 region to interact with m4-1BBL with minimal additional contacts from CRD1 and CRD3 (Figure 4B). CRD4 of m4-1BB is not in contact with the ligand and appears disordered in one of the two receptors. In the complex, m4-1BBL uses residues from the AB’, CD and GH loops to engage with 4- 1BB, while none of the residues from the β-strands contribute to the binding interface (Figure 4C). Moreover, the DE loop of m4-1BBL is remote and opposite to the site of interaction while in other TNF-TNFR complexes, residues of this region energetically favor the receptor binding. Detailed inspection revealed that hydrophobic and polar contacts are formed throughout the interface and play a major role in the complex formation while electrostatic interactions are observed to a lesser extent. The binding interface is divided into three regions (Figure 4C). Region 1 contains a relatively small contact area in which, Arg 38 of m4-1BB forms a salt bridge with Asp 170 of m4-1BBL while Phe 36 of m4-1BB forms vdW interaction with Gly 171 of m4-1BBL (Table 2). Region 2 consists of an elongated contact area, where the A1 module of CRD2 of m4-1BB grips the AB’ loop and the B2 module holds the CD and GH loops of m4-1BBL. Specifically, Asn 60, Ser 50 and Asn 82 of m4-1BB use their side chains to interact with the ligand residues Gly 173, Asn 155 and Tyr 291. Furthermore, m4-1BBL residues Asp 282, Asp 286 all interact with Tyr 70 of m4-1BB via hydrogen bonding, while Phe 210 forms an aromatic π-π stacking interaction (Table 2). At region 3, the A2 module of the CRD3 domain is placed nearly parallel to the CD loop of m4-1BBL while no sign of contact was observed between its A1 module and the ligand molecule. Residues involved in the interaction network at this region are Phe 91, Leu 94, Arg 100 and Cys 101 of m4-1BB and Pro 208, Thr 209, Phe 210 and Thr 211 of m4-1BBL.

**Table 2.** Interactions between m4-1BBL and m4-1BB.

| **m4-1BBL** | **m4-1BB (CRD1)** | **Polar contacts** |
| --- | --- | --- |
| Asp 170 (OD2) | Arg 38 (NH1) | 3.4 (Salt bridge) |
| Asp 170 (OD1) | Arg 38 (NH2) | 2.9 (Salt bridge) |
| **m4-1BB (CRD2)** |  |  |
| Gly 171 (O) | Cys 61 (N) | 2.7 |
| Gly 173 (N) | Cys 61 (O) | 2.8 |
| Gly 173 (O) | Asn 60 (ND2) | 3.0 |
| Tyr 291 (OH) | Asn 82 (ND2) | 2.8 |
| Tyr 291 (OH) | Ile 63 (O) | 2.6 |
| Asn 155 (ND2) | Ser 50 (OG) | 3.1 |
| Asp 282 (OD2) | Tyr 70 (OH) | 2.8 |
| **m4-1BB (CRD3)** |  |  |
| Thr 209 (O) | Cys 101 (N) | 2.8 |
| Pro 208 (O) | Arg 100 (NH2) | 2.8 |
| Thr 211 (N) | Cys 101 (O) | 3.5 |
| **m4-1BBL** | **m4-1BB** | **Non-polar contacts** |
| Gly 171 | Phe 36 | 3.7 |
| Ala 172 | Pro 48 | 4.0 |
| Ala 172 | Thr 51 (CG2) | 4.0 |
| His 167 | Ile 63 | 3.7 |
| Tyr 291 | Arg 65 (CB) | 3.7 |
| Ala 157 | Val 66 | 4.3 |
| Thr 209 | Phe 71 | 4.2 |
| Phe 210 | Tyr 70 | 3.9 |
| Phe 210 | Phe 91 | 3.5 |
| Thr 211 | Leu 94 | 4.0 |

### Receptor induced conformational changes in m4-1BBL

Structural comparison of free and receptor-bound m4-1BBL revealed significant restructuring of the ligand conformation that can be attributed to receptor binding. The major structural adaptations involve mainly the loop regions (Asn 155 - Leu 159 and Ser 168 - Tyr 176 of AA’ loop, Pro 208 - Thr 211 of CD loop and the Ser 290 - Asn 293 of GH loop) that are involved in direct interaction with the receptor (Figure 5A). All of these loops undergo 3 ~ 4 Å movements that alter the position of critical residues interacting with the receptor. Though not present at the binding interface, the DE and EF loops that were disordered in the free m4-1BBL become ordered upon m4-1BB binding. In the receptor-bound m4-1BBL, the EF loop acquires a proper α-helix like structure in both protomers because of the allosteric effect induced by reordering of the adjacent CD loop residues that were in contact with the receptor (Figure 5B). Not only the individual protomers undergo structural modifications, but their relative orientation and position with respect to each other change (RMSD of 2.4 Å over the entire protomer Cα atoms) (Figure 5B). Superposition of both free and receptor bound m4-1BBL onto one protomer reveals that while the overall structure of the protomers, with the exception of the CD and EF loops, is highly similar, protomer B of the receptor-bound m4-1BBL dimer is tilted by ~ 7 Å towards the dimer axis. This change in the dimeric orientation brings the F strands of both protomers nearby, resulting in the formation of novel inter-subunit polar contacts at the middle region of the dimer interface that were missing in the unbound m4-1BBL dimer (Figure 5C and Table 3). Consequently, in the receptor-bound m4-1BBL, both protomers buried a total surface area of ~ 1960 Å^2^ compared to 1600 Å^2^ in the free m4-1BBL dimer.

**FIGURE 5.**
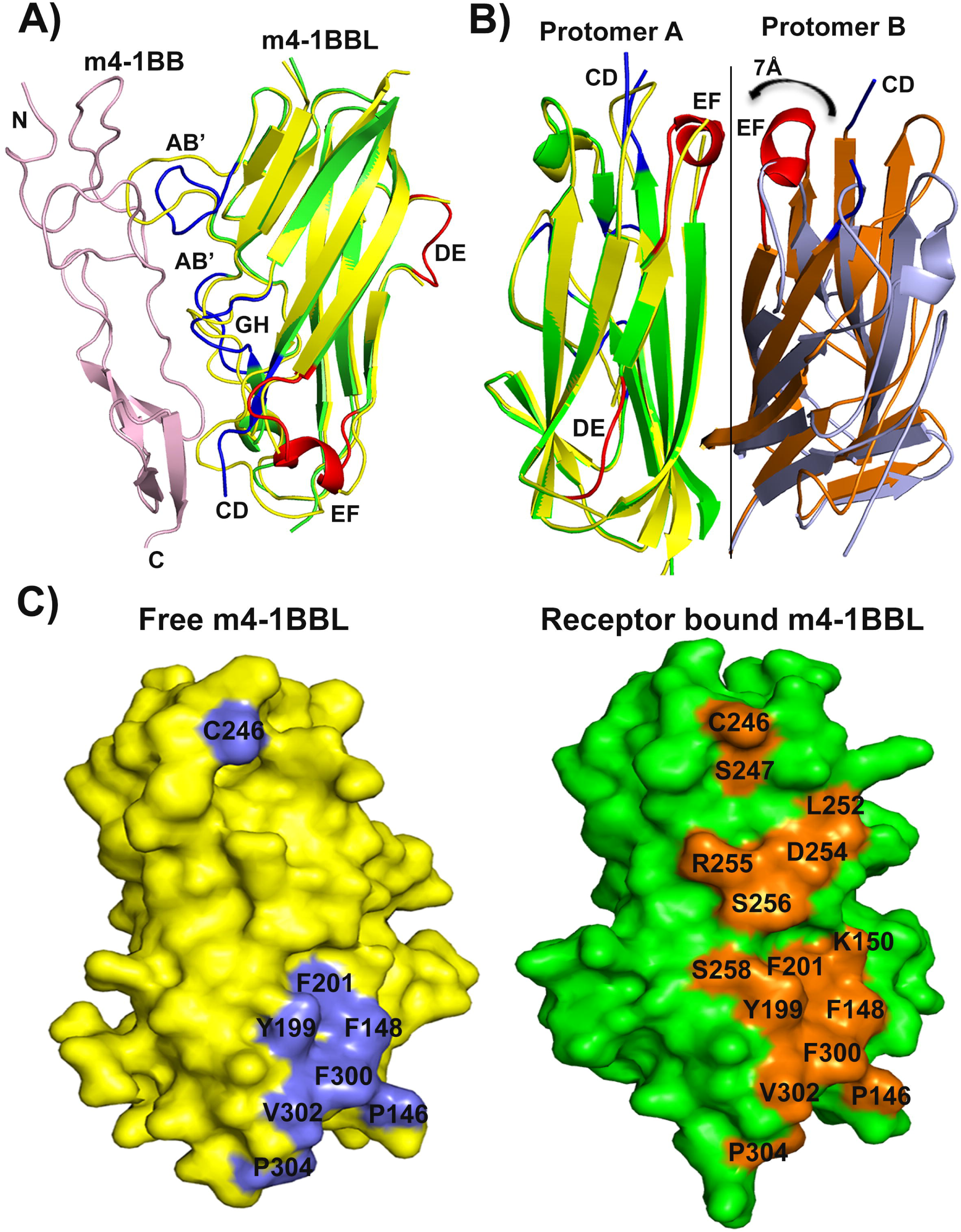
Receptor induced conformational changes of m4-1BBL. A) Superposition of free m4-1BBL monomer (yellow cartoon) with receptor bound m4-1BBL (green cartoon) showing select conformational changes in the loops (blue) which are in contact with receptor. The m4-1BB is shown in light pink color with its N and C-terminal ends marked. Structural ordering in the DE and EF loops of m4-1BBL are highlighted in red. B) Superposition of receptor bound m4-1BBL dimer with free m4-1BBL dimer discloses the proper alignment of protomer A and the deviation in the orientation of protomer B. The protomer A and B of receptor bound m4-1BBL are shown in green and orange and for free m4-1BBL are in yellow and blue. The EF loop that attains proper α-helix in both protomers induced by structural reordering of adjacent CD loop (blue) is highlighted in red. C) Binding foot print of protomer B on protomer A revealing the dimerization interface of free m4-1BBL versus receptor bound m4-1BBL. Residues participating in the dimeric interface of free m4-1BBL are colored blue and for receptor bound m4-1BBL are colored orange.

**Table 3.** Dimerization interface of receptor bound m4-1BBL dimer.

| **Protomer A**<br>**(m4-1BBL)** | **Protomer B**<br>**(m4-1BBL)** | **Polar contacts** |
| --- | --- | --- |
| Cys 246 (SG) | Cys 246 (SG) | 2.0 |
| Lys 150 (NZ) | Tyr 199 (OH) | 2.9 |
| Lys 150 (NZ) | Ser 258 (OG) | 3.3 |
| Ser 256 (N) | Asp 254 (OD1) | 3.7 |
| Ser 256 (OG) | Ser 256 (OG) | 2.6 |
| Tyr 199 (OH) | Lys 150 (NZ) | 2.8 |
| Cys 246 (SG) | Ser 247 (N) | 3.5 |
| Cys 246 (O) | Cys 246 (SG) | 3.4 |
| Leu 252 (O) | Arg 255 (NH2) | 2.5 |
| Asp 254 (OD2) | Ser 256 (N) | 3.1 |
| Ser 258 (OG) | Lys 150 (NZ) | 2.7 |
| Arg 255 (NE) | Asp 254 (OD2) | 3.1 |

### Binding characterization of m4-1BB/4- 1BBL complex

To determine the minimal binding requirements between m4-1BB and its ligand, we chose six residues of m4-1BB (Arg 38, Ser 50, Asn 60, Tyr 70, Asn 82 and Phe 91) that were spread over the 3 interaction regions for alanine mutagenesis and measured their binding affinity towards m4-1BBL using surface plasmon resonance studies. All of these variants are properly assembled as confirmed by size exclusion chromatography. Of these variants, the N82A and F91A resulted in undetectable binding up to 2 μM in SPR studies. Substitutions at Arg 38 had a moderate effect by decreasing the affinity ~5 fold (K_D_ =1.2 nM) while the affinity of N60A, S50A and Y70A were essentially unchanged towards binding to m4-1BBL (Figure 6). The three mutations that did affect the affinity of receptor for the ligand are dispersed over the m4-1BB/4-1BBL interface. Arg 38 is the only residue in CRD1 region that forms a salt bridge with the AB’ loop of the ligand (Figure 4C). Similarly, Asn 82 of CRD2 employs its side chain amide group to form a hydrogen bond with the OH group of Tyr 291 (GH loop) that protrudes deeply towards the interior of CRD2. Phe 91 of CRD3 forms a hydrophobic interaction (antiparallel π-π stacking) with Phe 210 (CD loop). Surprisingly, the π-π stacking interaction of Tyr 70 of CRD2 with Phe 210 (CD loop) appeared to be less important in binding compared to Phe91, since the affinity was essentially unchanged in the Y70A mutant. Our mutational analysis further confirmed that in contrast to other conventional TNF/TNFR complexes in which the DE loop of the ligand is most important for receptor binding, the m4-1BB/4-1BBL interaction is not concentrated in one location but spread out in at least two regions i.e. CD and GH loops of m4-1BBL.

**FIGURE 6.**
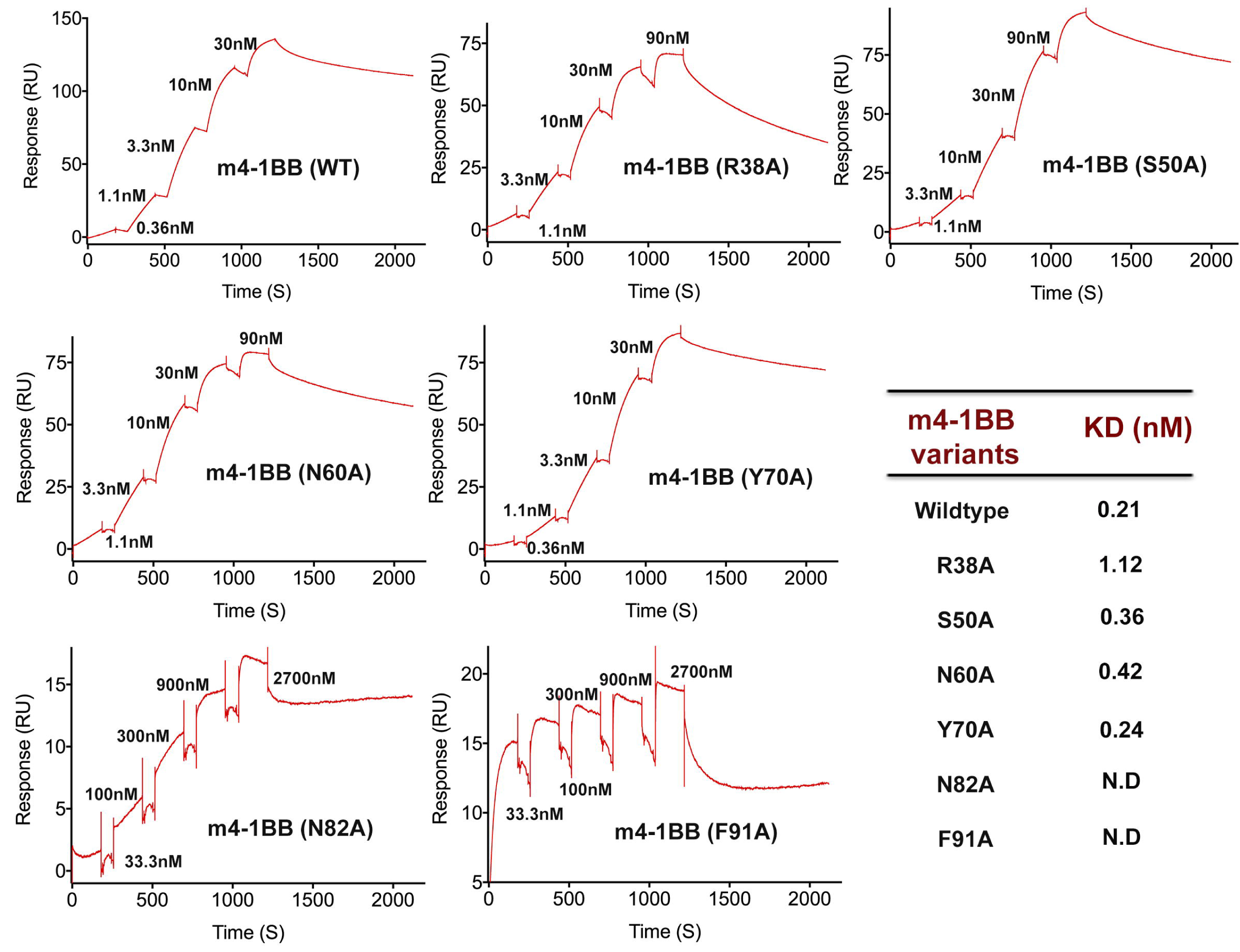
SPR analysis for the binding of various mutants of m4-1BB with m4-1BBL. Single cycle kinetics to measure the interaction of m4-1BBL with immobilized m4-1BB-Fc variants on anti Fc capture chip. SPR sensorgrams (red) were fitted by 1:1 binding model to measure the binding affinity between m4-1BB variants and m4-1BBL.

### Comparison to h4-1BB/4-1BBL complex

A number of TNFSF/TNFRSF exhibit cross-species interactions between the mouse and human molecules, in which the mouse ligand can interact with the human receptor with comparable binding affinity and the human ligand can interact with the corresponding mouse receptor. In contrast, 4-1BB/4-1BBL interactions are largely species specific. While h4-1BB does not bind to m4-1BBL, m4-1BB binds to h4-1BBL with greatly reduced affinity compared to that of h4-1BB (23). To understand the greatly restricted cross-species interactions, we compared the overall architecture of the respective 4-1BB/4-1BBL complexes (Figure 7). The major difference arises in the oligomeric assembly of the ligand, due to human 4-1BBL forming a hexameric functional unit and m4-1BBL forming a tetrameric signaling unit. Additionally, in the h4-1BB/h4-1BBL complex, the trimer is arranged in such a way that the binding site for each h4-1BB is formed by two adjacent protomers that provide a combined binding site, while m4-1BB only has a single binding site for a single m4-1BBL protomer.

**FIGURE 7.**
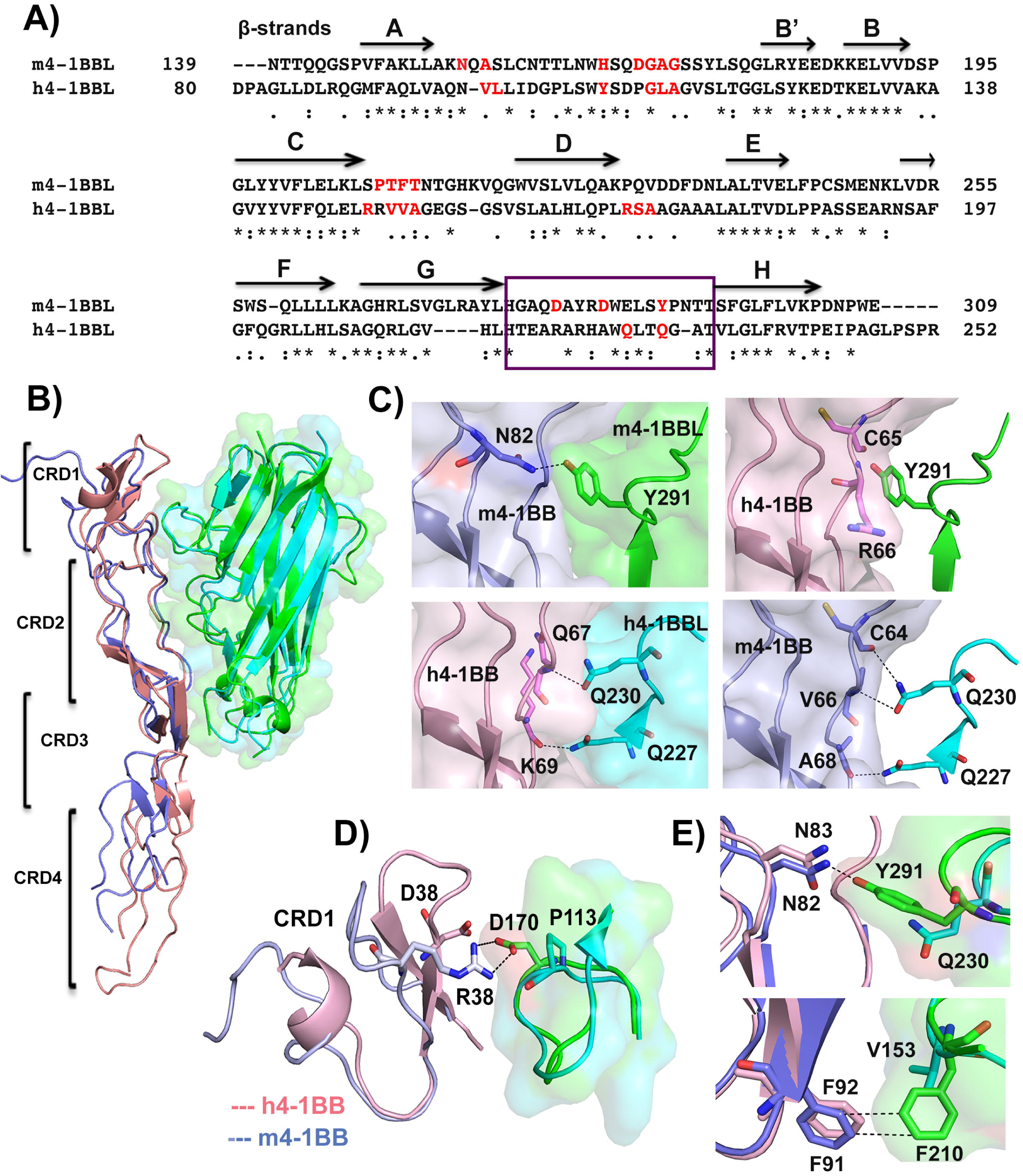
Comparison of 4-1BB/4-1BBL complex in human and mouse. A) Sequence alignment of THD regions of 4-1BBL of mouse and human. β-strands present in m4-1BBL are labeled. The residues involved in binding interface of the complex are colored red and the residues of GH loop that define species selectivity for 4-1BBL are boxed. B) Structural superposition of 4-1BB/4-1BBL complex in human and mouse by aligning the structurally equivalent β strands of the ligand. 4-1BBL of human and mouse are shown as cartoon with transparent surface in cyan and green color respectively. 4-1BB of human and mouse is represented as cartoon in light pink and light blue color. The four CRD regions of 4- 1BB are labeled. C) Interactions between GH loop residues and CRD2 region of m4-1BBL (green) and m4-1BB (blue) (top left); h4-1BBL (cyan) and h4-1BB (pink) (bottom left). The residues of h4-1BB that cause steric clash with Y291 of m4-1BBL are shown as pink sticks and the residues of m4-1BB that could bind to h4-1BBL are represented in blue sticks. D) Structural deviation between CRD1 region of m4-1BB (light blue) and h4-1BB (Light pink). The Arg 38 of m4-1BB CRD1 that makes salt bridge contact with Asp 170 of m4-1BBL (green) are shown as sticks. E) Interactions between N82 of m4-1BB and Y291 of m4-1BBL; F91 of m4-1BB with F210 of m4-1BBL. The N83 and F92 of h4-1BB (pink sticks) have no interacting partners in h4-1BBL (Cyan sticks). All the interactions are shown as black dashed lines.

A comprehensive evaluation between mouse and human 4-1BBL and 4-1BB identify substantial differences in their amino acid composition that correlate with their species specificity. The interface of the h4-1BB/h4-1BBL complex is mostly hydrophobic with few polar contacts that are mediated by main chain carbonyl and amide groups of polar residues. Conversely, the interface of m4-1BB/m4-1BBL complex is mostly polar and involves several residues engaging their side chains to form the complex (Figure 7A and Figure S2). Comparison of the binding interface in both complexes shows that both 4-1BB ligands have similar receptor binding areas confined to surface exposed AA’/AB’, CD and GH loops, however, their receptor interacting residues are not identical (Figure 7A and 7B). Apart from this, additional backbone interactions involving DE loop residues of adjacent protomer that are located in h4-1BB/4-1BBL complex are missing in the mouse complex. Sequence alignment revealed that amongst all receptor-binding loops, the receptor binding residues of GH loop are more divergent between human and mouse. Structural superimposition of both complexes demonstrates that the presence of a tyrosine residue (Y291) in the GH loop of m4-1BBL changes its conformation and displaces it by ~5 Å towards the receptor side compared to the GH loop of h4-1BBL. This conformation allows the Tyr 291 to protrude towards the interior of the B2 module of CRD2 thereby it makes hydrogen-bonding interaction with the side chain atoms of Asn 82 of m4-1BB (Figure 7C. In contrast, the GH loop of h4-1BBL is shorter and possesses two key receptor interacting residues Gln 227 and Gln 230 that make hydrogen-bonding contacts with main chain carbonyl and amide groups of Lys 69 and Gln 67 present at the surface of B2 module of CRD2 (Figure 7C). The N82A mutant of m4-1BB resulted in weak/undetectable binding for m4-1BBL and the Q227A and Q230A mutants of h4-1BBL decreased binding affinity by 80-fold (21) suggesting that GH loop interactions are energetically critical for the formation of 4-1BB/4-1BBL complexes in both mouse and human. When the structures of receptor complexes of h4-1BBL and m4-1BBL are superimposed by aligning the structurally equivalent β strands of the ligand, the GH loop residues Gln 227 and Gln 230 of h4-1BBL can retain binding with the CRD2 region of m4-1BB. On the other hand, because of the longer GH loop of m4-1BBL, its Tyr 291 sterically clashes with the backbone atoms of Cys 65 and Arg 66 of h4-1BB (Figure 7C, right panel) preventing the interaction between m4-1BBL and h4-1BB.

Both human and mouse 4-1BB also use different CRDs to bind their cognate ligands. M4-1BB uses residues from CRD1, CRD2 and CRD3 to bind m4-1BBL and those contacts made by all three CRDs contribute to the binding affinity. In contrast, in the h4-1BB/4-1BBL complex, residues from CRD2 and CRD3 are involved in the binding interface and the interactions from CRD2 residues actively contribute towards binding to h4-1BBL (Figure S2A). Although both receptors share ~60% sequence identity with analogous topology, superposition of m4-1BB with h4-1BB (both in complex with ligand), by aligning their structurally similar CRD2 region, results in an rmsd value of ~1Å (Figure S2C). While the CRD2 and CRD3 region of h4-1BB and m4-1BB superpose well, the CRD1 region exhibits structural distortion in comparison to h4-1BB. The CRD1 region of m4-1BB arranges as an extended loop instead of canonical anti-parallel β-strands that are typically observed in h4-1BB or any other TNFR members. In the m4-1BB/4-1BBL complex, the flexible nature of this loop brings its Arg 38 residue close enough to form a salt bridge with Asp 170 of the ligand (Figure 7D). This interaction seems to be dynamically significant as the Arg 38A mutant of m4-1BB has a ~5 fold lower affinity to its ligand compared to wild type. H4-1BB contains an aspartic acid in place of Arg 38 and the shorter side chain of Asp 38 is unable to form any interaction with h4-1BBL. Besides, most of the ligand-interacting residues of CRD2 and CRD3 of m4-1BB and h4-1BB are also not identical, however the two critical residues Asn 82 and Phe 91 that are essential for binding of m4-1BB to m4-1BBL are conserved in h4-1BB. Nonetheless, they do not make similar interactions in the h4-1BB/4-1BBL complex as they lack the partner residues in h4-1BBL (Figure 7E). Hence, from the comparison of both complexes, we can postulate that not only the unique organization of 4-1BBL, but select structural features at its GH loop region, and distinctive mode of 4-1BB binding in both human and mouse, restrict the species selectivity and define their specificity of interaction.

## DISCUSSION

In this study, we report the crystal structure of m4-1BBL alone as well as in complex with its cognate receptor m4-1BB. Rather than clustering into non-covalently associated bell-shaped homotrimers, m4-1BBL exhibits a dimeric quaternary structure in which a covalent inter-molecular disulfide linkage connects both protomers. This is unique, as the majority of TNF ligands possess intra-molecular disulfide bond-connecting cysteine residues from EF and CD loops of the same protomer (13) whereas m4-1BBL lacks this extra cysteine on its CD loop. Additionally, to avoid steric clashes with the longer CD and GH loops, the EF loop of m4-1BBL allosterically changes its conformation upon m4-1BB binding. This distinct orientation moves the EF loop of both protomers away from the monomer-monomer interface (Figure S3A and S3B). Further, the absence of conserved hydrophobicity on the ‘F’ strand of m4-1BBL (F199 of h4-1BBL replaced by Ser 256 in m4-1BBL) explains the scarcity of major inter-subunit contacts that would drive the assembly into a trimer.

In comparison to other TNF family members, human TRAIL also possesses a free cysteine on its EF loop similar to m4-1BBL (17,24). However, TRAIL’s GH loop is shorter, and hence the EF loop does not experience any conformational adjustment. In addition, in the trimeric interface, Tyr 237 and Gln 205 of this EF loop make hydrogen bonding contacts between adjacent protomers and the tyrosine residue on the ‘F’ strand communicates with neighboring units, resulting in proper aligning of individual subunits as a conventional homotrimer (Figure S3C). On account of this, we predicted that humanization of m4-1BBL via replacement of Cys 246 of the EF loop and Ser 256 of the ‘F’ strand with the human sequence would result in homotrimerization, and indeed it resulted in the appearance of differential oligomers in solution. Additionally, we visualized that reordering of the EF loop to a α-helix like structure in the receptor-bound m4-1BBL induces physical alterations in the relative orientation of its individual protomers. Of note, these results imply that any structural adaptations induced in either the EF loop or F strand region merely reorganize the whole monomer-monomer interface rather than assembling it as trimers. Based on this, we hypothesized that along with Cys 246 and Ser 256, the amino acid variations between m4-1BBL versus other TNF ligands might also be responsible for its unique behavior. Sequence alignment of 4-1BBL from various species revealed that the three critical residues (Cys 246, Ser 256 for dimerization, and Tyr 291 for receptor binding) present in m4-1BBL are conserved only in rodents and in the course of evolution, 4-1BBL has been acquired trimeric features in primates and other species (Figure S3D). The only other TNF ligand that exhibits a similar dimeric behavior is mGITRL, which contains a domain swapped interface in its C-terminal residues to stabilize the atypical dimer. Previous biochemical studies predicted that the dimeric mGITRL might also form a hetro-tetrameric complex with its receptor (25,26). HGITRL is also an extended trimer and lacks specificity towards mGITR (19), similar to trimeric h4-1BBL that exhibits very low affinity towards m4-1BB.

In general, most of the conventional TNF-TNFR complexes display a binding site formed by the surface exposed AA’, CD, DE and GH loops near the inter subunit cleft of two adjacent protomers. This allows each ligand trimer the opportunity to interact with three receptor molecules and also results in a binding mode where one receptor simultaneously binds to two adjacent ligand protomers. In this arrangement, the TRAF binding motifs of TNFR cytoplasmic tails come together in a close arrangement that favors recruitment of trimeric TRAF adaptors (13). The m4-1BB/m4-1BBL complex differs in that it forms an assembly with 2:2 stoichiometry, where one receptor exclusively interacts with a single subunit. In addition, the DE loop is not involved in receptor binding. This differences both in the stoichiometry of the receptor-ligand assembly and the details of their interaction suggests that m4-1BB could exhibit significant differences in how its signals are transduced compared to other conventional TNFR family molecules, as this atypical ligand/receptor assembly would not be favorable for a strong interaction with a trimeric TRAF. However, since CRD4 of m4-1BB can bind Galectin-9 via its N- and C-terminal carbohydrate binding domains (12,27), we proposed that this would bridge two m4-1BB molecules, leading to further oligomerization into tetrameric m4-1BBL complexes. This would then facilitate m4-1BB receptor oligomerization that could lead to strong TRAF recruitment and signaling similar to h4-1BB.

Two decades ago it was suggested that 4-1BBL and its receptor can participate in bidirectional signal transduction (28), and that 4- 1BBL reverse signaling has been found to transduce either positive or negative signals dependent on the cell expressing this ligand (29,30). Analogous to TNF, the cytoplasmic region of both human and mouse 4-1BBL contains a casein kinase 1 motif (SXX’SX where X can be any acidic amino acid), a site of phosphorylation proposed to be instrumental in reverse signaling (31). From our crystal structures we found that the distance between C-terminal ends of adjacent subunits in the m4-1BBL dimer and h4-1BBL trimer is ~ 26 Å and ~10 Å respectively, which is similar to that of TNF (8 Å) and GITRL (20 Å) that can also mediate reverse signaling. However, whether this results in a differential ability to signal or mediate different biological effects in human versus mouse cells is not clear. Previous studies reported some species variability in 4- 1BBL signaling (32). Cross-linking of h4-1BBL induced the maturation of monocytes, enhancing the expression of costimulatory molecules and secretion of cytokines like IL-12 and IFN-γ, but cross-linking of m4-1BBL did not activate murine monocytes in the same manner. It is possible that this difference might be explained by dimers versus trimers of 4-1BBL on the surface of the cells, although more studies would need to be performed to validate this hypothesis.

## EXPERIMENTAL PROCEDURES

### Design of m4-1BB and m4-1BBL constructs

For structural and binding studies, m4-1BBL was produced in Sf9 insect cells. Specifically, the cDNA encoding the ectodomain of m4-1BBL spanning the THD region along with additional C-terminal tail residues (amino acids 140-309) was cloned downstream of the gp67 secretion signal sequence into the baculovirus transfer vector pAcGP67A. An N-terminal hexa-histidine tag followed by a thrombin cleavage site (LVPRGS) was inserted upstream of m4-1BBL to assist its purification. For crystallization of the m4-1BBL - m4-1BB complex, the same construct of m4-1BBL without any purification tag was cloned into pAcGP67A vector. In parallel, the ectodomain of m4-1BB including all four cysteine rich domains (CRD 1-4; amino acids 24-160) was also cloned independently into separate pAcGP67A vector with a C-terminal hexa-histidine tag for co-expression. For SPR binding studies, m4-1BB was cloned into a modified mammalian expression vector pCR 3.1 downstream of the HA signal sequence and upstream of the Fc domain of human IgG1 and expressed in mammalian HEK 293T cells. The precise sequence for all the clones was confirmed by DNA sequencing.

### Generation of m4-1BBL and m4-1BB mutants

To generate single mutation variants of m4-1BBL and m4-1BB, site-directed mutagenesis was performed using Quick Change II site directed mutagenesis Kits (Stratagene, La Jolla, CA, USA). Single alanine mutations were made in the m4-1BB-Fc fusion protein at residues R38, S50, N60, Y70, N82 and F91. In the m4-1BBL construct, the cysteine at position 246 was substituted with serine (C246S mutant) and the serine at position 256 was replaced with phenylalanine residue (S256F mutant). In parallel, a double mutant of m4-1BBL carrying both of these mutations was also generated. The mutants were purified with a Qiagen mini prep kit and the presence of a mutation was verified by DNA sequencing. All the mutants of m4-1BB-Fc were expressed in mammalian HEK 293T cells and the mutants of m4-1BBL were expressed in Sf9 insect cells.

### Preparation of recombinant baculovirus

The baculovirus transfer vector pAcGp67A containing either wild type or mutant of m4-1BBL expression constructs having an N-terminal hexa-histidine tag were independently transfected into BacPAK6DNA under sterile conditions. To increase the efficiency, the transfection was performed in serum free media using Bacfectin reagent according to the manufacturer’s protocol. To obtain recombinant virus, the transfection mixture was prepared by gently mixing 1 μg of recombinant DNA, 100 ng of BacPAK6DNA and 5 μl of Bacfectin reagent that was filled up to 100 μl with sterile medium. As a control, another transfection mix without the BacPAK6DNA was also prepared. The experimental and the control transfection mix were incubated for 15 minutes in the dark and then added separately to the seeded 2 × 10^6^ healthy dividing *Spodoptera frugiperda* (Sf) 9 cells while gently swirling the T-25 flask and then incubated at 27°C in a medium containing 50 U/ml penicillin and 50 μg/ml streptomycin. After 7 days, the transfected virus with a multiplicity of infection below 1 (MOI<1) was collected by centrifugation at 1000×g, which then subsequently used for two rounds of virus amplification to achieve the virus titer having MOI=1. For protein production, high titer recombinant virus stock having MOI value ranging from 3 to 5 was added to individual 1 L cultures seeded with 2000 × 10^6^ Sf9 cells and the protein was expressed at 27°C by shaking at 145 rpm for 72-84 hrs. The supernatant containing the desired protein was collected by centrifugation for 10 mins at 1000×g.

### Expression of m4-1BB/4-1BBL complex

The extracellular region of m4-1BB was co-transfected with m4-1BBL in Sf9 insect cells using BacPAK6DNA. The recombinant virus stock for m4-1BB/4-1BBL complex was prepared similar to individual viral stocks. To achieve equal protein synthesis, the transfection mixture was prepared by mixing equal concentrations (2μg) of m4-1BB and m4-1BBL with 0.5 μg of BacPAK6DNA and 5μl of Bacfectin reagent in a total volume of 100 μl, transfected, and the protein was expressed as reported above.

### Protein purification of m4-1BBL and m4-1BB/4-1BBL complex from insect cells

For protein purification, the Sf9 cell supernatant containing the desired protein was further centrifuged at high speed to remove additional cell debris. The final supernatant was concentrated to ~ 300 ml while exchanging the buffer against 1X PBS (Phosphate buffered saline) by tangential flow filtration using 10 kDa molecular weight cut-off membranes (PALL). The individual native/mutants of m4-1BBL or m4-1BB/4-1BBL complex were purified by Ni^2+^ ion affinity chromatography. Briefly, 5 ml Ni-NTA resin was added to the concentrated supernatant and gently stirred overnight at 4°C. Later, the Ni beads were collected, washed with 20mM imidazole and the His-tagged fusion proteins were eluted with 250 mM imidazole (in 50 mM Tris HCl, 300 mM NaCl, pH 8.0) buffer. The proteins were further purified by size exclusion chromatography using Superdex S200 column in 50mM HEPES pH 7.5 and 150mM NaCl buffer. For crystallographic studies, the N-terminal his-tag was removed from m4-1BBL by thrombin cleavage using 5 units of bovine thrombin per mg of protein at 25°C. After 8 hours, the thrombin was inactivated by treatment with protease inhibitor PMSF and subsequently removed by size exclusion chromatography on a Superdex S200 column. Fractions containing cleaved wild-type or mutant versions of m4-1BBL (in 50mM HEPES pH 7.5 and 150mM NaCl buffer) were pooled and concentrated to ~ 10 mg/ml for crystallization. For structural studies of the m4-1BB/4-1BBL complex, the co-purified ligand/receptor complex was concentrated to 7 mg/ml and subsequently crystallized.

### Expression and purification of wild-type and mutant verions of m4-1BB from mammalian HEK293T cells

The protein expression and purification of m4-1BB-Fc fusion proteins in mammalian expression system has been reported previously (12). Briefly, the expression constructs of native/mutants of mouse 4-1BB-Fc were transiently transfected into mammalian HEK 293T cells using standard calcium phosphate transfection. After 3.5 days of protein expression at 37°C under 5% CO_2_, the supernatant containing secreted m4-1BB-Fc protein was collected and buffer exchanged against 1 X PBS. The wild-type and mutant proteins were purified by affinity chromatography using Protein A column followed by size exclusion chromatography (Superdex S200 column) in 50 mM HEPES and 150 mM NaCl buffer. The peak fractions were pooled, concentrated and stored at −80°C.

### Crystallization of m4-1BBL and m4-1BB/4-1BBL complexes

Initial crystallization trials for m4-1BBL and the m4-1BB/4-1BBL complex were performed in a 96-well format using a nano-liter dispensing liquid handling robot (Phenix, Art Robbins Ltd.). Over 600 different commercially available crystallization screens (JCSG core+, JCSG core 1-4 screens, Sigma) were tested by sitting drop vapor diffusion method at both 4°C and 22°C. Optimization of crystals was carried out in hanging drop by equilibrating 1.2 μl of protein and 0.8 μl of reservoir solution. Crystals of m4-1BBL grew in approximately 15 days in a well solution consisting of 2.4 M ammonium sulfate and 0.1 M MES pH 6.0. Crystals of m4-1BB/4-1BBL appeared in 2 days in various drops having PEG 4000 as a common precipitant. Among 20 crystallization conditions, the reservoir solution consisting of 0.2 M Ammonium chloride and 20% PEG 4000 generated high quality-diffraction crystals. Prior to crystal diffraction, all crystals were cryoprotected by immersing in mother liquor containing 20% glycerol (for m4-1BBL) or in a mixture of paratone oil and paraffin oil in 1:1 ratio (for m4-1BB/4-1BBL crystals).

### Data collection and refinement

Native X-ray diffraction data for all crystals were collected at Stanford Synchrotron Radiation Light Source beamline 9-2 at a wavelength of 0.97 Å and at 100 K temperature. Data images were collected with 0.15-degree oscillation and 1 - 2 sec exposure time for different crystals. The data images were indexed, integrated and scaled in HKL 2000 package (33) to an overall resolution of 2.5 Å for m4-1BBL. For m4-1BB/4-1BBL complex, multi crystal dataset was generated by merging two individual native datasets in AUTOPROC (34) and STARANISO (35) to an overall resolution of 2.62 Å. For structure solution of m4-1BBL, we have used the recently reported h4-1BBL structure (PDB 6D3N) as a search model for molecular replacement method in PHASER-MR (36,37). Concurrently, the position of m4-1BB/4-1BBL complex in the asymmetric unit was also determined by similar method using the structures of our current m4-1BBL and previously reported m4-1BB (PDB 5WI8) as search models. The models were further refined with PHENIX/REFMAC (38) and BUSTER (39) with tight non-crystallographic symmetry restraints. The surface exposed loops of m4-1BBL and the CRD3 region of m4-1BB in the m4-1BB/4-1BBL complex were built in the Fo-Fc electron density map gradually by cycles of iterative manual model building with program COOT (40,41) and ARP/wARP (42) function as part of the CCP4 suite (43,44). At last phase of refinement, N-glycans and water molecules were added. The final structure of m4-1BBL and m4-1BB/4-1BBL complex were refined to residual factors R/Rfree= 18.8/24.1 and 20.4/23.8 respectively. In the final model, both structures have more than 98% residues in the favored region of Ramachandran plot. The data collection and refinement statistics are summarized in Table 1. All figures were made in PyMOL (45).

### SEC-MALS analysis

A miniDAWN TREOS multi-angle light scattering detector, with three angles (43.6°, 90° and 136.4°) detectors and a 658.9 nm laser beam, (Wyatt Technology, Santa Barbara, CA) in combination with the Optilab T-rEX refractometer (Wyatt Technology) were used in-line with the Agilent Technologies 1200 Series liquid chromatography system (Agilent Technologies, Santa Clara, CA) for size exclusion chromatography analysis. Samples (10 micrograms) were injected onto the size exclusion chromatography analytical column, AdvanceBio SEC 300Å, 7.8 x 300 mm, 2.7 μm column (Agilent Technologies) with 0.2M phosphate, pH 7 as the mobile phase at a flow rate of 0.5 mL/min for a duration of 40 minutes at 25°C. Detection was done using a DAD detector at 214 and 280 nm signal. Data collection and SEC analysis were performed using ChemStation software. Data collection and dynamic light scattering analysis were performed using ASTRA 6 software (Wyatt Technology).

### Surface Plasmon Resonance binding kinetics

The binding affinity between m4-1BBL and various mutants of m4-1BB was determined by surface plasmon resonance studies using a BIACORE T200 instrument at 293K. m4-1BB-Fc variants at a concentration of 20 μg/ml were captured on CM5 sensor chip (GE health care) immobilized with anti-human IgG (Fc) to get ~ 300 response units (active surface). A reference sensor surface captured with free Fc was used as a negative control to subtract non-specific binding. The binding and kinetic experiments were performed in assay buffer (HBS-EP) composed of 10mM HEPES, 150 mM NaCl, 3 mM EDTA and 0.05% Tween 20 and all ligands and analytes were diluted in this buffer prior to the experiment. The kinetic constants for the binding of m4-1BBL with variants of m4-1BB were determined by single cycle kinetics method. Increasing concentrations of m4-1BBL as an analyte were injected over active and reference surfaces at a flow rate of 30 μl/min and the rate of association was recorded for 180-240 s followed by a 100 s gap in between each injection and a final dissociation was monitored for an additional 900 s. The kinetics for the interaction of m4-1BBL with immobilized m4-1BB variants was monitored in real time and expressed with a sensorgram reporting magnitude of response in relative units. The data was analyzed using the Biacore T200 Evaluation software 2.0 (GE Healthcare) using a kinetic model describing 1:1 binding between analyte and ligand to calculate the equilibrium dissociation constant (*K*_D_ = *k*_d_/*k*_a_) by non-linear fitting.

## Acknowledgements

The authors thank the Stanford Synchrotron Lightsource (SSRL) for access to remote data collection and the SSRL beamline scientists for support.

## Author contributions

AB conducted all of the biochemical and structural experiments, analyzed the results and wrote the manuscript. TD assisted in data collection and phasing. GD performed SEC-MALS studies. MC conceived the overall project with DMZ and edited the paper. DMZ conceived the experiments, supervised the overall project and wrote the manuscript.

## Footnotes

This project has been funded in part with federal funds from the National Institute of Allergy and Infectious Diseases, National Institutes of Health, grant <u>AI110929</u> (MC and DMZ).

Use of the Stanford Synchrotron Radiation Lightsource, SLAC National Accelerator Laboratory, is supported by the U.S. Department of Energy, Office of Science, Office of Basic Energy Sciences under Contract No. DE-AC02-76SF00515. The SSRL Structural Molecular Biology Program is supported by the DOE Office of Biological and Environmental Research, and by the National Institutes of Health, National Institute of General Medical Sciences (including P41GM103393). The contents of this publication are solely the responsibility of the authors and do not necessarily represent the official views of NIGMS or NIH.

